# Uncovering the Vital Role of Lipid Droplets in Coronavirus Replication: A Novel Insight Arising from the Biological Activity of 2-Bromopalmitate

**DOI:** 10.1101/2023.06.29.547086

**Authors:** Dongxiao Liu, Ruth Cruz-cosme, Julian Leibowitz, Yong Wu, Qiyi Tang

## Abstract

The emergence of viral infections with global impact highlights the urgent need for broad-spectrum antivirals. In this study, we evaluated the effect of palmitoylation inhibitors [2-bromopalmitate (2-BP), cerulenin, and 2-fluoro palmitic acid (2-FPA)] and the enhancer palmostatin B on the replication of human coronaviruses (hCoV-229E, hCoV-Oc43) and murine hepatitis virus (MHV-A59) at non-cytotoxic concentrations. The results demonstrated that 2-BP strongly suppressed MHV-A59 replication, while cerulenin and 2-FPA only moderately inhibited viral replication. Palmostatin B significantly enhanced viral replication. Notably, 2-BP exhibited superior efficacy. Interestingly, palmostatin B failed to rescue the inhibitory effects of 2-BP but effectively rescued cerulenin and 2-FPA, suggesting additional biological activities of 2-BP beyond palmitoylation inhibition. Furthermore, we discovered that 2-BP specifically disrupted lipid droplets (LDs), and this LD disruption was correlated with viral replication inhibition. Based on our findings, we conclude that the inhibitory effects of 2-BP on viral replication primarily stem from LD disruption rather than palmitoylation inhibition. Therefore, we revealed the crucial role of LDs in the viral replication. Our study provides insights into the development of wide-spectrum antiviral strategies.

## INTRODUCTION

Viral diseases represent a significant portion of global infectious diseases and are responsible for a substantial number of annual deaths worldwide. According to estimates by the World Health Organization (WHO), viral diseases contribute to approximately one-third of the total global deaths each year (1). Current antiviral development has predominantly focused on targeting viral processes or enzymes. This strategy aims to achieve potent antiviral effects while minimizing host-cell toxicity by avoiding cross-inhibition of host proteins (2). Despite the progress made in antiviral drug development, the lack of a wide-spectrum drug against diverse viruses remains a challenge. The emergence and reemergence of viral infections can occur unpredictably, necessitating the urgent development of broad-range antivirals. To address this, a new approach focuses on targeting host-cell pathways and enzymes that are essential for virus replication. Viruses depend on the host-cell environment for their replication, making virus-host interactions crucial during the disease process. By disrupting these virus-dependent host-cell pathways and enzymes, broad-spectrum antivirals can effectively inhibit viral replication across different viral strains. This paradigm shift offers promising opportunities for the development of effective antivirals capable of combating a wide range of viral infections.

The significance of fatty acid oxidation (FAO) in viral replication and pathogenesis has gained increasing recognition (3–7). Fatty acids obtained from the extracellular environment or cellular lipid storage, such as lipid droplets (LDs), serve as essential resources for FAO. The primary outcome of FAO is the production of ATP, which is crucial for viral replication (4). Notably, the inhibition of FAO using chemical compounds has demonstrated promising effects in suppressing cell growth and inducing apoptosis, highlighting its therapeutic potential against viral infections. LDs are ubiquitous organelles in mammalian cells, and their role in viral infection and replication remains a subject of debate. Some studies have reported an increase in LD numbers and abundance during infection with viruses such as hepatitis C virus (HCV) and rotavirus, suggesting that LD induction is necessary for viral replication (8, 9). However, it has also been observed that Dengue virus (DENV) infection triggers lipophagy, leading to a decrease in LDs (10). Hence, the exact role of LDs in enhancing or suppressing viral replication requires further investigation and clarification.

In our study, we investigated the inhibitory effects of 2-bromopalmitate (2-BP) on coronaviruses, specifically Murine hepatitis virus strain A59 (MHV-A59), hCoV-229E, and hCoV-OC43. MHV-A59 is a beta-coronavirus belonging to the same genus as SARS-CoV-2, sharing similar replication mechanisms and cycles (11–13). MHV-A59 infection in mice serves as a valuable small animal model for studying beta-coronavirus pathogenesis and antiviral drug development. While previous studies have reported the inhibitory effects of 2-BP on MHV replication (14), the underlying mechanisms remained unclear. In our study, we utilized the MHV-A59 infection system in 17CL-1 cells to investigate the impact of different inhibitors and enhancers of S-palmitoylation on viral infection, gene expression, and replication. Our findings revealed that 2-BP exhibited significantly stronger repression of viral replication compared to the other two palmitoylation inhibitors, cerulenin and 2-fluoro palmitic acid (2-FPA). Interestingly, the S-palmitoylation enhancer, palmostatin B, which targets acyl protein thioesterase 1 (APT1) and 2 (APT2) in cells, significantly promoted viral replication. A noteworthy discovery was that 2-BP, but not cerulenin or 2-FPA, caused a pronounced disruption of lipid droplets (LDs), suggesting that the effects of 2-BP on viral replication might be attributed to its influence on LDs. Importantly, we observed that 2-BP degraded LDs and inhibited viral replication at a concentration that had no significant impact on palmitoylation. These findings highlight the potential role of LDs in viral replication and suggest that 2-BP’s inhibitory effects on viral replication may be mediated through its disruption of LDs rather than its direct impact on palmitoylation. This provides valuable insights into the mechanisms underlying the antiviral activity of 2-BP and opens avenues for further research on the interplay between LDs and viral infections.

## RESULTS

### 2-BP exhibits significantly stronger inhibitory effects on MHV-A59 replication compared to other palmitoylation inhibitors, namely cerulenin and 2-FPA

In this study, we investigated the effects of palmitoylation inhibitors and enhancers (Figure S1A and S1B) on the replication of Murine hepatitis virus strain A59 (MHV-A59), a beta-coronavirus, in 17CL-1 cells (15, 16). We used an MHV-A59 strain tagged with an enhanced Green Fluorescent Protein (eGFP) reporter gene, allowing us to visualize and photograph viral infection and replication under a fluorescence microscope (17).

The cells were infected with MHV-A59 at different multiplicities of infection (MOIs) and treated with either 2-bromopalmitate (2-BP), cerulenin, 2-fluoro palmitic acid (2-FPA), or palmostatin B. The concentrations of the drugs were determined to be non-cytotoxic based on the MTT assay performed earlier (Figure S1C-S1F). A no-drug group (the same concentration of DMSO) was used as a control (Figure 1A1-1A3).

**Figure 1.**
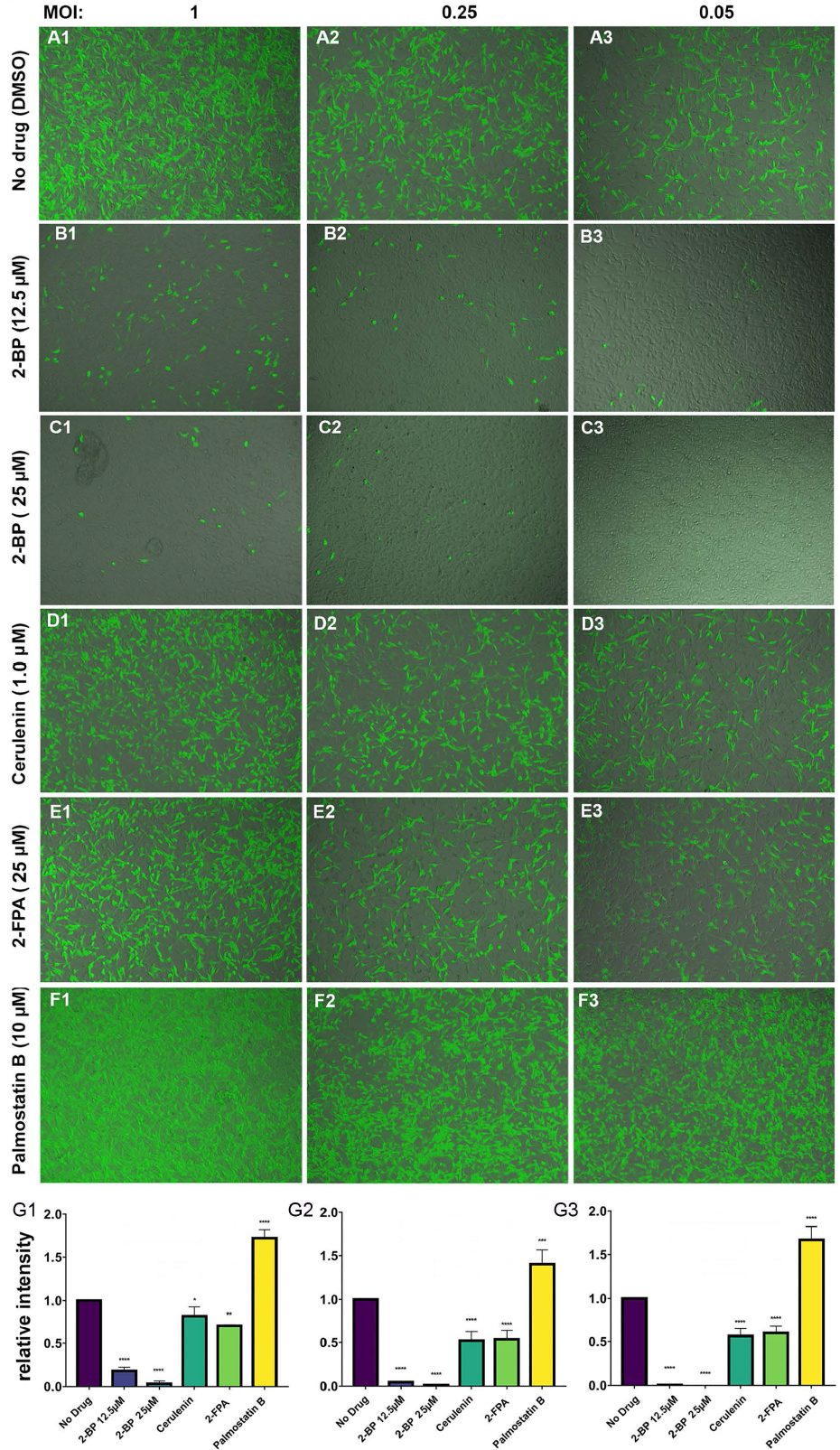
Fluorescence (GFP) to show the effects of palmitoylation inhibitors or an enhancer on the infection and replication of MHV-A59 in 17CL-1 cells. MHV-A59 was added together with the drug at a concentration as indicated for 24 hours. The images were taken under a fluorescent microscope at a 10 x len under the same exposure time. **A1-A3.** No drug; **B1-B3.** 12.5 µM of 2-BP; **C1-C3.** 25 µM of 2-BP; **D1-D3.** 1.0 µM of cerulenin; **E1-E3.** 25 µM of 2-FPA; **F1-F3.** 10 µM of palmostatin B. **G1-G3.** Three independent experiments were performed and the average GFP intensities were normalized to those of DMSO groups. The averaged numbers were compared to “no drug” group and shown as mean ± standard error (SE). *P<0.005, **P<0.001, ***P<0.0005, ****P<0.0001.

Fluorescent microscopy images of infected cells were captured 24 hours post-infection (hpi) to assess viral replication, and the representative images were shown in Figure 1. The results showed that 2-BP strongly suppressed MHV-A59 replication at both concentrations tested (12.5 µM and 25 µM). At 25 µM, 2-BP almost completely blocked replication even at a low MOI of 0.05. In contrast, cerulenin and 2-FPA only moderately inhibited MHV-A59 replication compared to the control group. Palmostatin B, the palmitoylation enhancer, significantly enhanced viral replication at a concentration of 10 µM. The average GFP intensities from three independent experiments were quantified and normalized to the DMSO control group, as shown in Figure 1G1 to 1G3. Our results showed that 2-BP has a significantly stronger inhibitory effect on MHV-A59 replication than the other two palmitoylation inhibitors.

Although the immunofluorescence assays (IFA) as shown in Figure 1 clearly demonstrated the inhibitory effects of 2-BP and enhancive effects of palmostatin B on MHV-A59 replication in 17CL-1 cells, viral protein production, and viral growth curve assays are necessary to be examined to show the viral replication and gene expressions. To that end, immunofluorescence assays and western blot assays were performed. MHV-A59-infected cells were treated with the drugs, and whole-cell lysates were collected at different times post-infection. Western blot assays using anti-GFP, anti-N, and anti-Tubulin antibodies were conducted to assess viral protein production (Figure 2A). The results demonstrated that 2-BP strongly suppressed viral protein production compared to the control group, while cerulenin and 2-FPA showed lesser inhibitory effects. Palmostatin B increased the production of GFP and N proteins significantly. The results from three independent experiments were statistically averaged and analyzed as shown in Figure 2B for GFP and Figure 2C for N protein. We demonstrated that 2-BP has much stronger inhibitory effects on viral protein production than the other two palmitoylation inhibitors, cerulenin, and 2-FPA.

**Figure 2.**
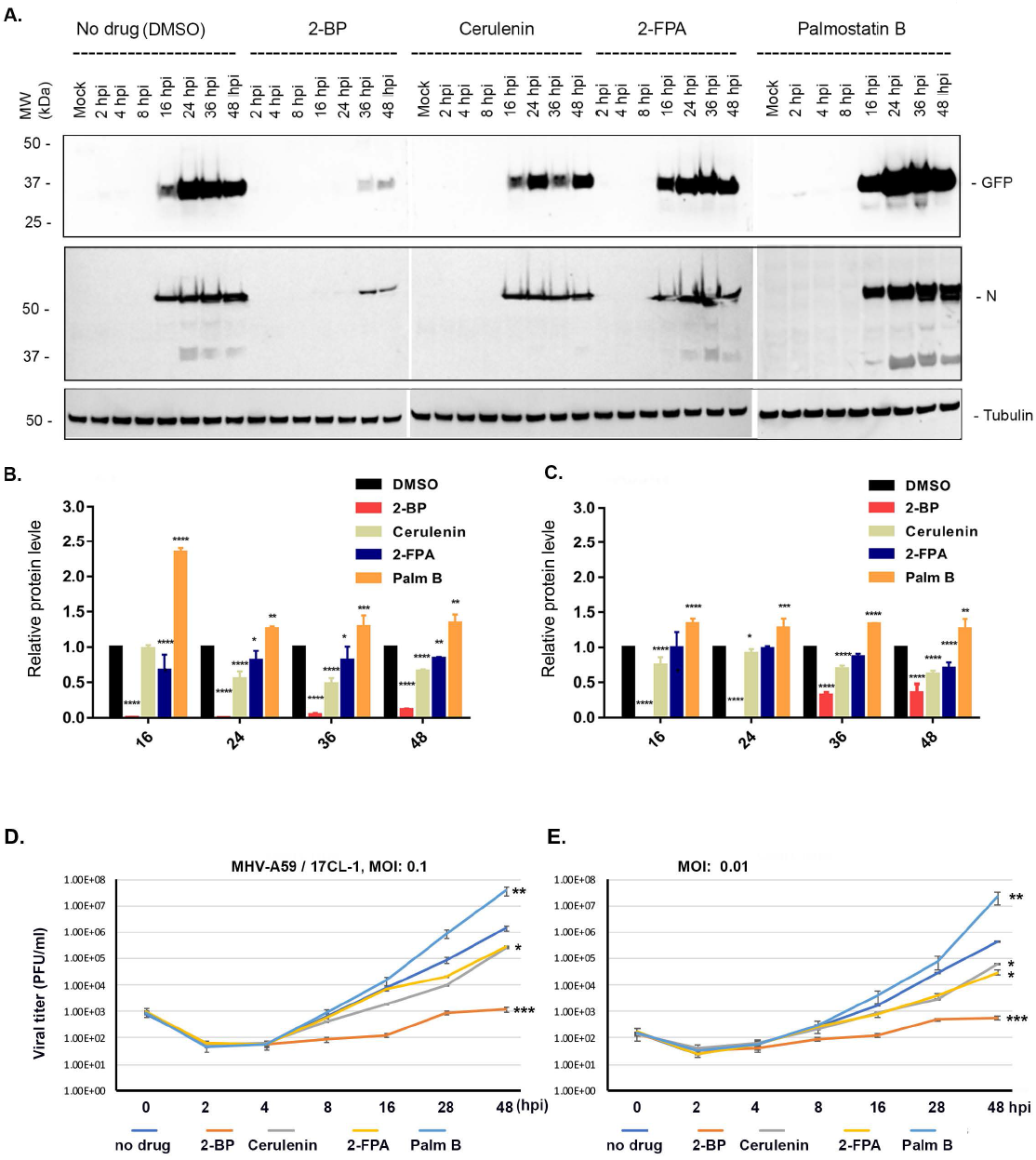
Palmitoylation inhibitors or an enhancer on the protein production and replication of MHV-A59 in 17CL-1 cells. **A** Western blot assay. The 17CL-1 cells were infected or non-infected with MHV-A59 at an MOI of 0.5 at the same time when were untreated or treated with the drug. The cells were collected at the time post-infection as indicated and lysed in lamellae buffer for western blot assay using the antibodies as shown on the right side. The experiments were performed independently three times and the representative ones were shown. **B and C.** The results from three different western blot experiments were statistically averaged and analyzed for GFP and N protein. 2-BP has much stronger inhibitory effects on viral protein production than the other two palmitoylation inhibitors, cerulenin, and 2-FPA. **D and E.** growth curve assay. MHV-A59 was added onto 17CL-1 cells at an MOI of 0.1 (**D**) or 0.01 (**E**). The cells and the medium were collected, and viruses were released by thawing and freezing for 3 times. The viral titers were determined by PFU assay. The experiments were performed independently 3 times and the average numbers were shown as mean ± standard error (SE). *p<0.005; **p<0.001; ***p<0.0001

To determine the growth curves of MHV-A59 in the presence or absence of drugs, a plaque-forming unit (PFU) assay was performed. Cells were infected with MHV-A59 at an MOI of 0.1 (Figure 2D) or 0.01 (Figure 2E) without or with a drug as indicated. Supernatants were collected at different time points, and viral titers were measured. Consistent with the results of the western blot assays, 2-BP strongly inhibited viral replication, while cerulenin and 2-FPA had minor inhibitory effects. Palmostatin B significantly increased viral replication. Overall, these findings indicate that palmitoylation inhibitors have suppressive effects on MHV-A59 infection and replication, while a palmitoylation enhancer promotes viral replication. The results suggest that protein palmitoylation plays a role in viral infection and replication. Notably, 2-BP exhibited stronger inhibitory effects on MHV-A59 replication than the other inhibitors, indicating the possibility of additional mechanisms of action for 2-BP.

### Palmostatin B fails to rescue the inhibitory effects of 2-BP, while it successfully mitigates the inhibitory effects of cerulenin and 2-FPA on MHV-A59 replication

Next, we investigated the ability of Palmostatin B, an inhibitor of APT-1 and APT-2 enzymes that enhance palmitoylation, to rescue the inhibitory effects of palmitoylation inhibitors on MHV-A59 replication. Our hypothesis posited that if 2-BP had additional mechanisms for inhibiting viral replication, its inhibitory effects could either be partially or not at all rescued by Palmostatin B. To test this, we conducted rescue experiments by combining Palmostatin B with each of the palmitoylation inhibitors.

As shown in Figure 3A-3C, the combination of 2-BP and Palmostatin B failed to rescue the inhibitory effects of 2-BP on MHV-A59 protein production (Figure 3A-3B) and viral replication (Figure 3C). There were no significant differences observed in protein levels and viral titers between the 2-BP alone group and the 2-BP plus Palmostatin B group. However, Palmostatin B significantly rescued the suppressive effects of cerulenin (Figure 3D-3F) and 2-FPA (Figure 3G-3I) on MHV-A59 protein production and viral replication. Our rescue experiments provide evidence supporting our hypothesis that 2-BP may employ additional mechanisms for inhibiting MHV-A59 replication, which is not effectively rescued by Palmostatin B.

**Figure 3.**
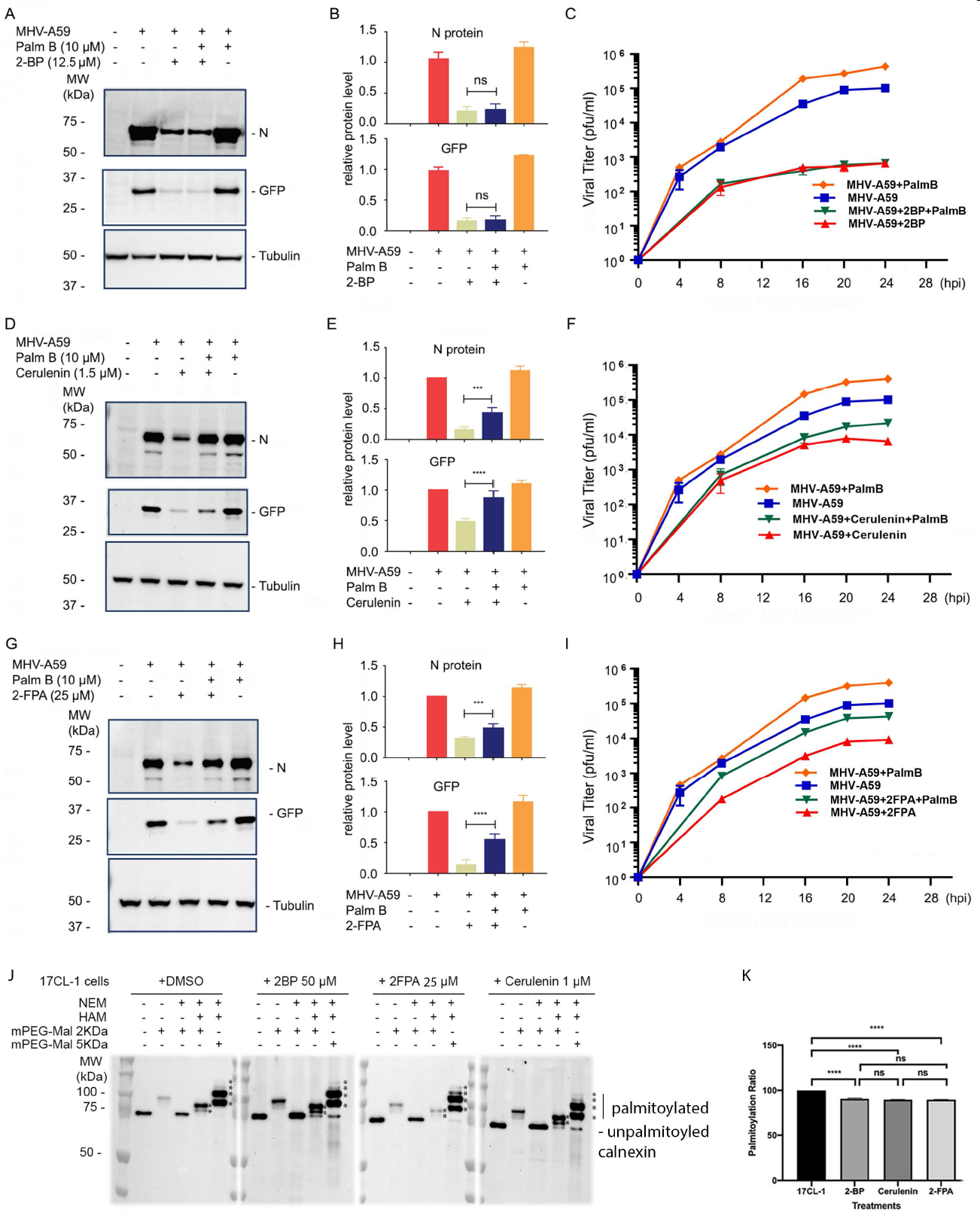
Rescuing the inhibitory effects of the palmitoylation inhibitors on MHV-A59 replication by palmostatin B. **A-C** The combination of 2-BP with palmostatin B. MHV-A59 was used to infect 17CL-1 cells in the presence of DMSO, Palm B (palmostatin B), 2-BP, or Palm B and 2-BP. Western blot was performed at 24 hpi (**A**) and the results from three different western blot experiments were statistically averaged and analyzed for GFP and N protein (**B**). MHV-A59 viral growth curve was determined as shown in **C**. **D-F.** The combination of cerulenin with palmostatin B. The same as A-C, but cerulenin was used instead of-BP. **G-I.** The combination of 2-FPA with palmostatin B. The same as A-C, but 2-FPA was used instead of 2-BP. **J & K.** APE assay. 17CL-1 cells were treated with DMSO, 2-BP, 2-FPA, or Cerulenin for 24 hours. The whole cell lysates were applied for APE assay as described in the “Materials and Methods”. The APE samples were used for western blot assay for calnexin that has been usually used as palmitoylation marker. The palmitoylated calnexin was labeled by asterisks. The palmitoylated bands were then compared to the unpalmitoylated band to calculate the palmitoylation rates as shown in K. The experiments were performed independently 3 times and the average numbers were shown as mean ± standard error (SE). *p<0.005; **p<0.001; ***p<0.0001.

We utilized the acyl-PEG exchange (APE) assay to compare the effects of different inhibitors (2-BP, cerulenin, and 2-FPA) on the palmitoylation of calnexin. Calnexin, an ER protein, is commonly used as a marker for palmitoylation assays (18, 19). 17CL-1 cells were treated with DMSO (control), 50 µM 2-BP, 1 µM cerulenin, or 25 µM 2-FPA for 24 hours. Cell lysate samples were subjected to NEM and HAM treatments, followed by incubation with PEG-2kDa or PEG-5kDa. NEM was used to prevent nonspecific labeling of PEG. In Figure 3J, the second lanes of each group show that PEG-2kDa nonspecifically binds to unblocked calnexin. The third lane demonstrates complete blockage of unspecific binding due to NEM treatment, revealing only unpalmitoylated calnexin. Slower-moving bands in lanes 4 and 5 indicate palmitoylated calnexin, with lane 4 showing PEG-2kDa labeling that was not well-separated to identify other palmitoylated calnexin bands (indicated by stars). The slower-moving bands in lane 5 represent PEG-5kDa-exchanged calnexin, displaying four palmitoylated calnexin bands (indicated by stars).

Using ImageJ, we compared the palmitoylated bands to the unpalmitoylated band in lane 5s to calculate the palmitoylation rate (Figure 3K). Interestingly, we observed no significant differences in calnexin palmitoylation among the inhibitor-treated groups, despite the significant inhibition of palmitoylation compared to the DMSO group. These findings provide further evidence that the inhibitory effect of 2-BP on MHV-A59 replication may involve mechanisms other than palmitoylation inhibition.

### Disruptive Effects of 2-BP on Lipid Droplets Revealed by Immunofluorescence Assay (IFA)

In our search for additional mechanisms by which 2-BP inhibits MHV-A59 replication, we conducted immunofluorescence assay (IFA) experiments to investigate the effects of 2-BP treatment on the distribution and intensity of various subcellular organelles, including the endoplasmic reticulum (ER), lysosomes, Golgi apparatus, mitochondria, late endosomes, and lipid droplets (LDs). The 17CL-1 cells were treated with 25 µM 2-BP for 24 hours and then stained using specific markers as shown in Figure 5: calnexin for ER (Figures S2A1 and S2A2), LAMP1 for lysosomes (Figures S2B1 and S2B2), Giantin for the Golgi apparatus (Figures S2C1 and S2C2), CoxIV for mitochondria (Figure S2D1 and S2D2), Rab7 for late endosomes (Figure S2E1 and S2E2), and BODIPY for LDs (Figure S2F1 and S2F2). The upper panels depict cellular images after 2-BP treatment, while the lower panels show images of cells without 2-BP treatment for comparison.

**Figure 5.**
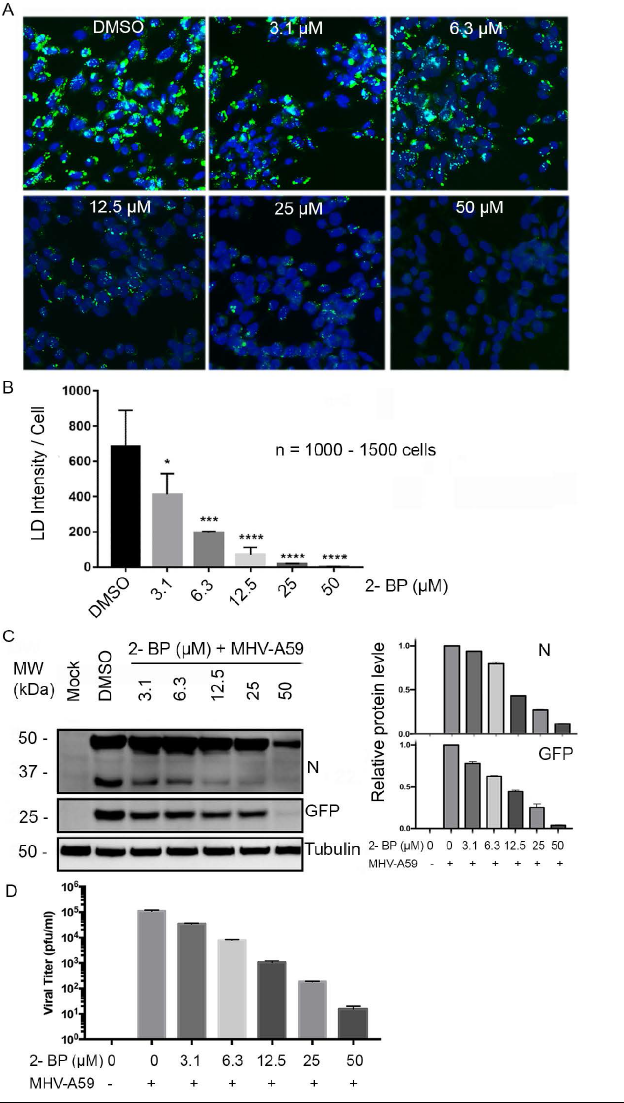
2-BP’s disruptive effect on LD is correlated to its inhibitory effects on MHV-A59 in 17CL-1 cells. **A &B** 2-BP’s disruption of LD is dose-dependent. 17CL-1 cells were treated with 2-BP at different concentrations as indicated for 24 hours and LDs were stained using BODIPY. Representative images were shown and the LD intensity per cell was quantified by ImageJ (n=1000-1500 cells). The bars indicate means ± SEM. *P < 0.05; **P < 0.01; ***P < 0.001, ****P < 0.001 (one-way ANOVA). **C.** 17Cl-1 cells were infected with MHV-A59 at MOI of 0.1. Serial dilution of 2-BP was added to the infected cells. Samples were collected after 16hrs’ incubation and subjected to western blot. MHV-A59-GFP-N and GFP were detected. The result was subsequently analyzed by ImageJ. **D.** 17Cl-1 cells were infected with MHV-A59 at MOI of 0.05. Serial dilution of 2-BP was added to the infected cells. the medium supernatant samples were collected after 16hrs’ incubation and subjected to plaque assays. Three independent experiments were performed, and the numbers were statistically analyzed.

Our IFA results revealed no significant changes in the intensity and structure of the ER, lysosomes, Golgi apparatus, mitochondria, and late endosomes upon 2-BP treatment. However, it is evident that LDs were almost completely eliminated following 2-BP treatment.

Furthermore, we investigated whether other palmitoylation inhibitors could also lead to LD degradation. For this purpose, we treated 17CL-1 cells with DMSO (Figure 4A), 2-BP (Figure 4B), Cerulenin (Figure 4C), 2-FPA (Figure 4D), or Palmostatin B (Figure 4E) for 24 hours. After the treatment, the cells were incubated with BODIPY 500/510 for 30 minutes at 37°C, followed by washing with PBS and staining the nuclei using DAPI. The images were captured under the same conditions, including the exposure time and UV strength. Compared to the untreated cells (Figure 4A), Cerulenin, 2-FPA, or Palmostatin B did not significantly affect LD density. However, 2-BP clearly reduced the intensity of LDs. Therefore, we discovered a previously unreported activity for 2-BP, which depletes LDs in the cells.

**Figure 4.**
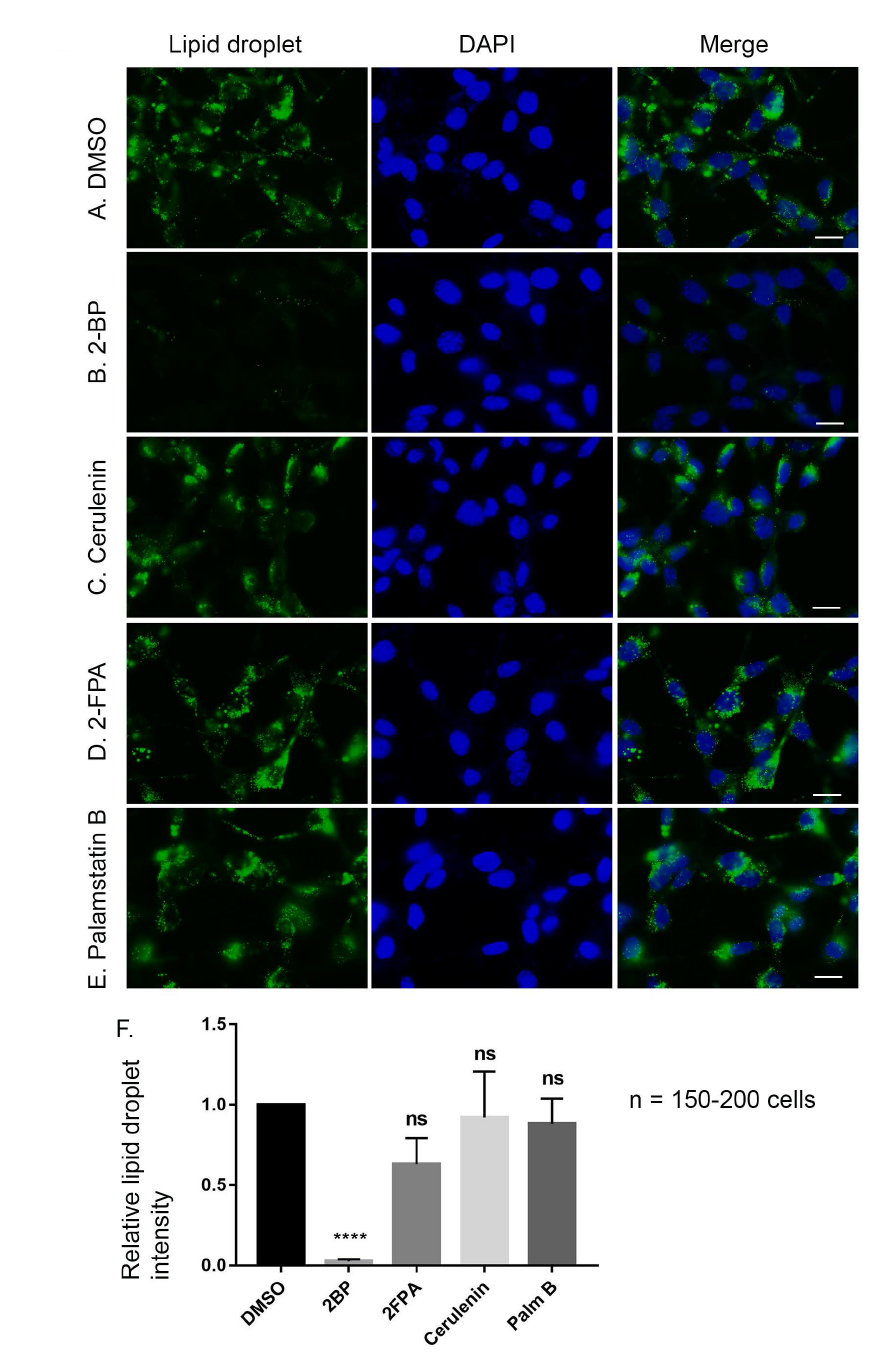
The effects of palmitoylation inhibitors or an enhancer on lipid droplets in 17CL-1 cells. The 17CL-1 cells were seeded on a coverslip and incubated for 24 hours. Then the cells were treated with No drug (DMSO, **A**), 25 µM of 2-BP (**B**); 1.0 µM of cerulenin (**C**); 25 µM of 2-FPA (**D**); or 10 µM of palmostatin B (**E**) for 24 hours. The cells were stained with BODIPY 500/510 for 30 min at 37°C. After being fixed with 1% paraformaldehyde for 10 min at room temperature, the cells were stained with DAPI for 1 min. The cells were visualized under a florescent microscope at a 40x len and the images were taken at the same exposure time and power. Scale bar: 10 μm. **F.** The averaged lipid droplet intensities were scanned from 150-200 cells by ImageJ and normalized to those of DMSO groups. _****_ p<0.0001, ns = not significant.

### A dose-dependent correlation between the inhibition of MHV-A59 by 2-BP and the depletion of LDs: Increasing the concentration of 2-BP leads to a more pronounced disruption of LDs and greater inhibition of MHV-A59 replication

Our aforementioned results demonstrated that 2-BP has a disruptive effect on LDs, which may relate to its inhibitory effects on MHV-A59. To demonstrate the correlation between 2-BP’s inhibition on MHV-A59 and its depletion of LDs, we set out to determine whether its inhibition on MHV-A59 and disruption of LDs are both dose-dependent. As shown in Figure 5A, we treated 17CL-1 cells with different concentrations of 2-BP as indicated for 24 hours. The LD (green) was visualized together with nuclei (blue) and images were shown in Figure 5A. The more concentrations of 2-BP treatment caused fewer LDs. We then scanned 100 to 1500 cells for each group by ImageJ program to quantify the intensity of LD. As can be seen in Figure 5B, the depletion of LDs by 2-BP is dose-dependent.

Next, we applied the same concentration series of 2-BP together with MHV-A59 to infect 17CL-1 cells for 24 hours. The whole cell lysates were collected at 24 hpi for western blot assays. As can be seen in Figure 5C, viral N protein, and GFP protein are both decreased in a dose-dependent manner. Then, we examined the inhibitory effects of the serially diluted 2-BP on viral replication. The same results were also observed for viral growth inhibition by 2-BP as shown in Figure 5D. Therefore, we suspect that 2-BP’s disruptive effects on LDs may be associated with its inhibitory effects on MHV-A59.

### LD depletion results in reduced viral protein production and viral growth

To investigate whether LDs would reappear in cells following LD depletion caused by 2-BP treatment, we conducted a time course experiment. 17CL-1 cells were treated with 50 µM 2-BP or DMSO for 24 hours, trypsinized, and then re-cultured on coverslips. LD staining was performed at various time points post-trypsinization (TIN). As shown in Figures 6A and 6B, the 2-BP-treated cells remained devoid of visible LDs for nearly 24 hours after TIN, while still maintaining healthy growth. The cell cultures after TIN no longer contained 2-BP or DMSO. Subsequently, the trypsinized cells were infected with MHV-A59 at an MOI of 0.1 at different time points post-TIN (6, 12, 24, 36, 42, and 54 hours post-TIN). Whole cell lysate samples were collected 24 hours post-infection (hpi), and viral protein levels were analyzed by western blot. Figure 9C demonstrates that infections occurring during the LD-depleted period exhibited significantly lower viral protein production compared to the DMSO control groups. Statistical analysis confirmed these significant differences, as indicated in Figure 6C.

**Figure 6.**
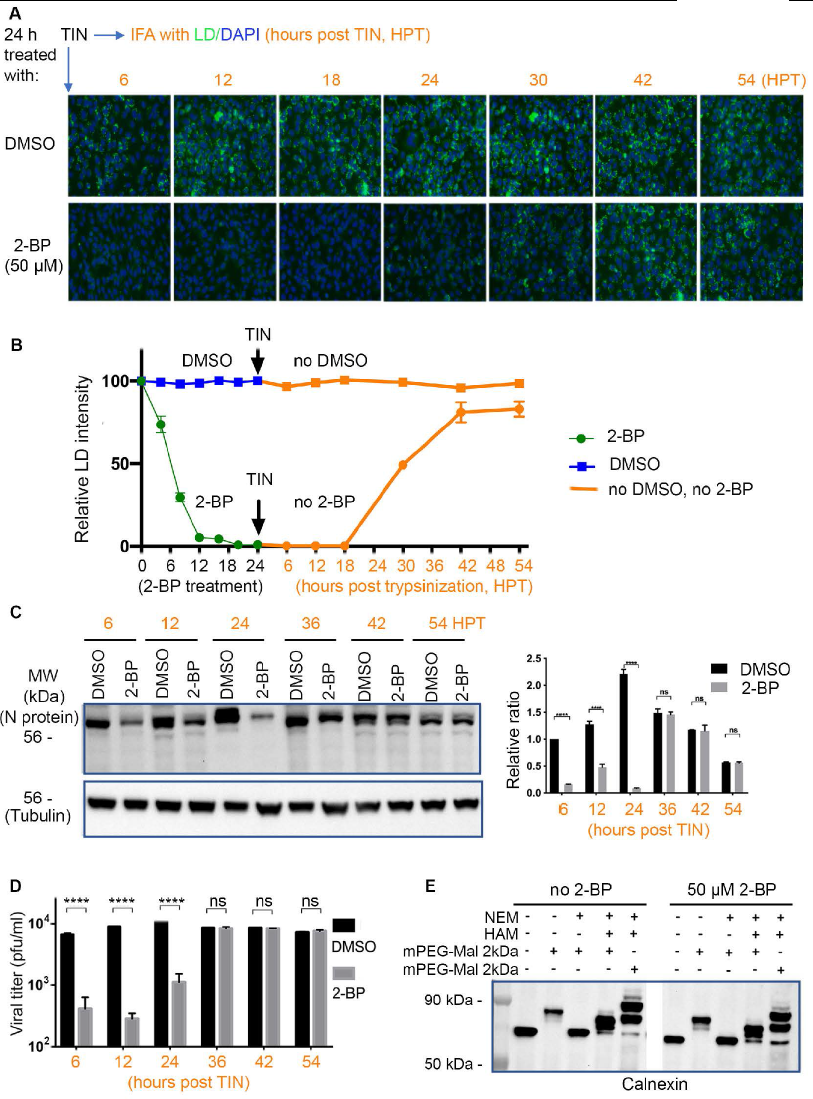
LD depletion results in reduced viral protein production and viral growth. **A** 17CL-1 cells were treated with 50 µM 2-BP (lower) or DMSO (upper) for 24 hours, trypsinized (TIN), and then re-cultured on coverslips. LD staining was performed at 6, 12, 18, 24, 30, 36, 42, 48, 54 hours post-trypsinization (TIN) and photographed under a 40 x len for LD and DAPI. **B.** More than 5 images were scanned every 6-hour post treatment or post-TIN.for each group. Average intensities were calculated by comparing to 0-hour post treatment. **C.** The trypsinized cells were infected with MHV-A59 at an MOI of 0.1 at 6, 12, 24, 36, 42, and 54 hours post-TIN. Whole cell lysate samples were collected 24 hours post-infection (hpi), and viral protein levels were analyzed by western blot for N protein (upper) and tubulin lower. **D.** The trypsinized cells were infected with MHV-A59 at an MOI of 0.01 at different time points post-TIN (6, 12, 18, 30, 42, and 54 hours post-TIN). Cells and medium were collected 24 hours post-infection (hpi), and viral particle formation was quantified using plaque assays. Statistical analysis indicated. **E.** The acyl-PEG exchange (APE) assay to know whether 2-BP at a concentration of 50 µM exerted inhibitory effects on palmitoylation. Cell lysate samples were treated with NEM or HAM, or both, followed by incubation with PEG-2kDa or PEG-5kDa. Slower-moving bands represent palmitoylated calnexin. ****P < 0.001.

A similar experiment was conducted to assess whether LD-depleted cells exhibited reduced viral growth. The trypsinized cells were infected with MHV-A59 at an MOI of 0.01 at different time points post-TIN (6, 12, 18, 30, 42, and 54 hours post-TIN). Cells and medium were collected 24 hours post-infection (hpi), and viral particle formation was quantified using plaque assays. As depicted in Figure 6D, infections occurring during the LD-depleted period showed significantly lower viral production compared to the DMSO control groups. Statistical analysis indicated no significant differences in viral protein production or viral particle formation in later time points post-TIN (30/36, 42, and 54 hours post-TIN) (Figure 6D).

Furthermore, we employed the acyl-PEG exchange (APE) assay to investigate whether 2-BP at a concentration of 50 µM exerted inhibitory effects on palmitoylation. As shown in Figure 6E, we used calnexin as a marker for palmitoylation assay as reported previously (18, 19). Figure 9E displays the results of the palmitoylation assay using calnexin as a marker, as previously reported. Cell lysate samples were treated with NEM or HAM, or both, followed by incubation with PEG-2kDa or PEG-5kDa. Slower-moving bands represent palmitoylated calnexin. As observed, there were no significant differences in the palmitoylation of calnexin between the control group and the 50 µM 2-BP-treated group. Thus, we provide further evidence that the inhibitory effect of 2-BP on MHV-A59 replication at a concentration of 50 µM is primarily attributed to LD depletion.

### 2-BP induces LD depletion in human cells and exhibits inhibitory effects on human coronaviral infection and replication

Finally, we investigated whether 2-BP could inhibit the replication of human coronaviruses. Initially, we examined the effects of 2-BP on hCoV-OC43, a human beta coronavirus, in HCT-8 cells, which are permissive to hCoV-OC43 infection (20). Treatment with 50 µM 2-BP (a non-toxic concentration, Figure S3) led to LD depletion in HCT-8 cells (Figure 7A). Subsequently, we evaluated the growth curve of hCoV-OC43 in HCT-8 cells with and without 2-BP treatment. As depicted in Figure 7B, 2-BP significantly reduced the growth of hCoV-OC43 in HCT-8 cells. To assess viral protein production, we treated hCoV-OC43-infected HCT-8 cells with various concentrations of 2-BP and detected the nucleocapsid protein (N) using western blot analysis. Figure 7C illustrates that 2-BP dose-dependently inhibited viral N protein levels. Furthermore, when 50 µM 2-BP was applied during hCoV-OC43 infection in HCT-8 cells, the production of viral protein was significantly suppressed compared to the DMSO-treated group (Figure 7D).

**Figure 7.**
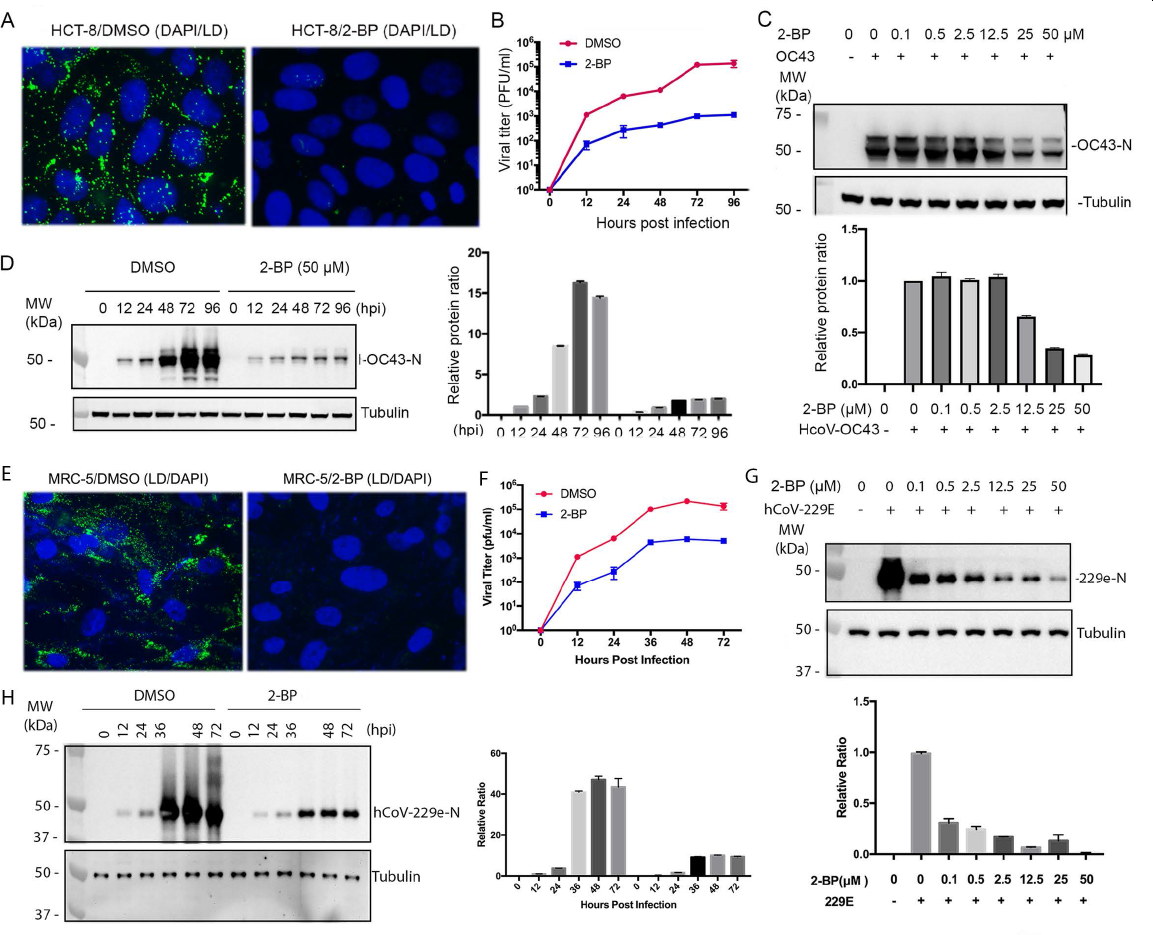
2-BP induces LD depletion in human HCT-8 cells and exhibits inhibitory effects on hCoV-OC43 and hCoV-229E infection and replication. **A** Treatment with 50 µM 2-BP led to LD depletion in HCT-8 cells. **B.** the growth curve of hCoV-OC43 in HCT-8 cells with and without 2-BP treatment. **C.** To assess viral protein production, we treated hCoV-OC43-infected HCT-8 cells with various concentrations of 2-BP and detected the nucleocapsid protein (N) using western blot analysis. **D.** 50 µM 2-BP was applied during hCoV-OC43 infection in HCT-8 cells, the production of viral protein was significantly suppressed compared to the DMSO-treated group. **E.** Treatment with 50 µM 2-BP led to LD depletion in MRC-5 cells. **F.** the growth curve of hCoV-229E in MRC-5 cells with and without 2-BP treatment. **G.** To assess viral protein production, we treated hCoV-229E-infected MRC-5 cells with various concentrations of 2-BP and detected the nucleocapsid protein (N) using western blot analysis. **H.** 50 µM 2-BP was applied during hCoV-229E infection in MRC-5 cells, the production of viral protein was significantly suppressed compared to the DMSO-treated group.

Since both hCoV-OC43 and MHV-A59 are beta-coronaviruses, we investigated whether 2-BP exhibited inhibitory effects on alpha-coronaviruses. We employed the hCoV-229E infection system in MRC-5 cells for this purpose. Similar to the findings in beta-coronaviruses, 2-BP at a non-toxic concentration (Figure S4) caused LD depletion in MRC-5 cells (Figure 7E). Plaque formation assays demonstrated that 2-BP reduced hCoV-229E growth in MRC-5 cells compared to the control group treated with DMSO (Figure 7F). Western blot analysis revealed that 2-BP dose-dependently inhibited hCoV-229E protein production in MRC-5 cells (Figure 7G). Additionally, the western blot experiments conducted during the infection time course showed strong suppression of viral protein production by 2-BP (Figure 7H), consistent with its inhibition of viral growth. Thus, 2-BP not only inhibits the replication and infection of beta-coronaviruses but also suppresses alpha-coronaviruses.

## DISCUSSION

Viral diseases have been posing disastrous consequences globally, so it is urgent to develop antivirals against emergent or reemergent viruses. Two main strategies, host-targeted antivirals (HTAs) and direct-acting antiviral agents (DAAs), have been employed in antiviral research. While DAAs have shown success in clinical settings by targeting viral proteins (21), their effectiveness is limited to specific viruses, and viral resistance often arises. Moreover, DAAs are not readily applicable to new viral infections until their characteristics are known. In contrast, HTAs have the potential to overcome the limitations of DAAs. Since all viruses rely on cellular machinery for replication, HTAs theoretically offer a wide-spectrum antiviral capability. This broad-spectrum approach becomes particularly valuable during emergent viral pandemics or epidemics. Our studies on the antiviral effects of 2-BP against MHV-A59, hCoV-OC43, and hCoV-229E serve as an example of developing a wide-spectrum antiviral. Given that 2-BP demonstrated antiviral effects on beta-coronaviruses such as MHV-A59 and hCoV-OC43, it is plausible that it may exhibit similar efficacy against other coronaviruses, including SARS-CoV-2.

Our study has yielded significant biological findings with implications for the field of antiviral research. Firstly, we have discovered a novel biological activity of 2-BP, demonstrating its ability to deplete LDs in various cell lines at non-toxic concentrations. This finding highlights a previously unknown effect of 2-BP. Secondly, contrary to its conventional role as a palmitoylation inhibitor, our results indicate that 2-BP can inhibit viral replication even at concentrations where its impact on palmitoylation is minimal. Instead, the depletion of LDs appears to be the primary mechanism responsible for the inhibitory effects of 2-BP on viral replication. Thirdly, the observed correlation between LD depletion and suppressed viral replication suggests a crucial role for LDs in viral replication processes. This finding further emphasizes the importance of LDs in the viral life cycle. Lastly, based on our study results, we propose the hypothesis that LDs serve as a source of energy for viral replication, and their depletion disrupts the energy supply, leading to the inhibition of viral replication. In summary, our research reveals that 2-BP disrupts LDs in various cells, even at concentrations where its effects on palmitoylation are minimal. This disruption of LDs is closely linked to the critical role of LDs in the replication of coronaviruses.

In our study, we employed a unique experimental model to demonstrate the significance of LDs in viral replication. We discovered that the depletion of LDs using 2-BP is a reversible process, with LDs returning within 24 hours in 17CL-1 cells. To investigate the impact of LDs on viral replication, we treated cells with 2-BP for 24 hours to deplete LDs, followed by trypsinization and reseeding in the absence of 2-BP. We observed that the cells lacked LDs at this time point, but 24 hours later, LDs reappeared. Crucially, we conducted virus infection experiments comparing cells with and without the return of LDs. The results revealed that when the virus infected cells during the period when LDs had not yet returned, viral replication was significantly lower compared to when LDs were present. This direct evidence strongly supports the importance of LDs in viral replication. Importantly, our investigation extended beyond beta coronaviruses and included an alphacoronavirus, hCoV-229E. By testing the effects of LD depletion on multiple types of coronaviruses, we demonstrated the potential of targeting LDs as a strategy for developing broad-spectrum antivirals against RNA viruses.

Indeed, the role of lipid droplets (LDs) in viral replication is a complex and dynamic area of research, and different studies have presented contrasting findings. Monson et al. reported that LDs are involved in defending against viral infections caused by Zika virus and HSV-1 (22). They proposed that LDs play a role in cellular innate immunity, suggesting an inhibitory effect on viral replication. This adds another dimension to the functions of LDs as cellular organelles. However, these findings are not consistent across all viral infections. For instance, there is a contradictory report indicating that Dengue virus (DENV) infection leads to a reduction in LDs (10). This discrepancy suggests that LDs could have dual effects on viral infection: inhibitory in some circumstances and enhancing in others. The contrasting results observed in different studies may arise from variations in experimental conditions, virus strains, cell types, or other factors. It highlights the complex interplay between LDs and viral infections, and further research is needed to fully elucidate the mechanisms underlying these dual effects. Understanding the context-dependent nature of LDs in viral infections is crucial for comprehensively deciphering their role and developing targeted antiviral strategies. It emphasizes the need for further investigation to unravel the intricate relationship between LDs and viral replication, considering the diverse outcomes observed in different viral infection models.

Two theories may coexist for LDs’ effects on viral replication. The first theory suggests that LDs play a crucial role in viral replication by providing a source of energy for viral gene expression and replication. Viruses can utilize the lipids stored in LDs, such as triglycerides and cholesterol esters, as a fuel source to generate free fatty acids (FAs) for mitochondrial energy production. This energy is necessary for efficient viral replication and gene expression. Additionally, LDs may facilitate the assembly and maturation of viral particles. Targeting the LD-virus interaction could be a promising strategy for developing antivirals, as disrupting LD metabolism could impair viral replication. The second theory proposes that LDs are involved in cellular innate immunity. LDs are known to play a role in the regulation of immune responses and the production of interferons. By degrading LDs, viruses may inhibit the LD-mediated production of interferons, allowing them to evade the host immune system. Consequently, targeting the LD-virus interaction could not only disrupt viral replication but also enhance the cellular innate immune response, leading to a more effective antiviral defense. Furthermore, LDs are not essential for the viability of most cells, making them attractive targets for therapeutic interventions. Anti-diabetic or anti-obesity drugs that reduce LDs have been developed and widely used clinically. Leveraging this knowledge, developing broad-spectrum antivirals by targeting the LD-virus interaction holds promise. In this context, 2-BP, which has demonstrated the ability to deplete LDs and inhibit viral replication, could serve as an initial compound for further investigations and the development of antiviral strategies. The biological and clinical significance of your present studies lies in shedding light on the intricate relationship between LDs and viral replication, opening up new avenues for the development of broad-spectrum antivirals targeting LD-virus interactions.

In summary, unlike other palmitoylation inhibitors such as cerulenin and 2-FPA, 2-BP not only inhibits protein palmitoylation but also depletes LDs from cells. This unique property sets 2-BP apart from other inhibitors and suggests that its inhibitory effects on viral replication may be mediated through LD depletion rather than solely through palmitoylation inhibition. The strong inhibitory effect of 2-BP on viral replication indicates that LD depletion plays a crucial role in restricting viral infection. This observation is particularly relevant for coronaviruses and potentially applicable to other RNA viruses as well. It is important to consider the possibility that 2-BP’s effects on viral replication, achieved through LD depletion, may involve the elimination of LDs’ defensive functions against viruses. LDs have been implicated in cellular innate immunity and their degradation by viruses may enable viral evasion of the host immune response. Based on these findings, future studies will focus on further understanding the mechanisms underlying LD involvement in viral replication. This will involve investigating the precise role of LDs in viral infection and exploring strategies to target the LD-virus interaction. Additionally, research will be conducted to evaluate the potential of 2-BP as a broad-spectrum antiviral agent and to assess its feasibility in developing effective antiviral strategies.

## MATERIALS AND METHODS

### Tissue culture and virus

Murine cell line, 17CL-1 (derived from NIH3T3) was obtained from BEI resources (NIH/NIAID, NR-53719), HCT-8 (ATCC, CCL-244) and MRC-5 (ATCC, CCL-171) cells are maintained in Dulbecco’s Modified Eagle’s medium (DMEM) supplemented with 10% fetal calf serum (FCS), penicillin (100 IU/ml)-streptomycin (100 ug/ml) and amphotericin B (2.5 ug/ml) (23). Murine coronavirus, MHV-A59 (murine hepatitis virus strain A59) was obtained from BEI resources (NIH/NIAID, NR-53716) that is a recombinant Murine Coronavirus MHV-A59 with the replacement of ORF4 by Enhanced Green Fluorescent Protein (eGFP) (17). Human coronaviruses, hCoV-OC43 (NIH/NIAID, NR-52725) and -229E (NIH/NIAID, NR-52726) were obtained from BEI resources. MHV-A59, hCoV-OC43 and -229E were amplified in 17CL-1, HCT-8 and MRC-5 cells respectively. They were purified, tittered and stored at −80°C.

### Chemical Reagents

2-Bromohexadecanoic acid (2-BP, also called 2-bromopalmitate, cat# sc-251714), 2-Fluoropalmitic acid (2-FPA, cat# Sc-202881), Cerulenin (sc-200827) were purchased from Santa Cruz Biotechnology Inc, and Palmostatin B (Cat# 178501) was purchased form Sigma. They were solved in DMSO and store at −20°C.

### Antibodies

The monoclonal antibody against MHV-A59 nucleocapsid (N) protein was obtained from BEI resources (NIH/NIAID, NR-45106). Rabbit antibodies against hCoV-OC43 N (40643-T62) and hCoV-229E N (40640-T62) were purchased from SinoBiological US Inc. (Wayne, PA). Mouse antibodies against Tubulin (4G1, sc-58666), late endosomes (Rab7, B-3, sc-376362), monoclonal against GFP (sc-9966), and LAMP1 (sc-18821) were purchased from Santa Cruz Biotechnology (Santa Cruz, CA). Rabbit anti-FLAG (PA1-984B) was purchased from Invitrogen (Carlsbad, CA). Rabbit antibodies against Giantin (ab80864), CoxIV (ab16056), and calnexin (ab22595), were purchased from Abcam (Boston, MA).

### Immunofluorescent Assay (IFA)

Cells grown on coverslips were fixed with 1% paraformaldehyde for 10 minutes (min) at room temperature and permeabilized in 0.2% Triton for 20 min on ice. Immunostaining was performed by sequential incubation with primary antibodies and Texas red (TR)-labeled secondary antibodies (Vector Laboratories, Burlingame, Calif.) for 30 min each (all solutions in PBS). Washing steps using PBS were performed after each incubation with paraformaldehyde, Triton, or antibodies, after antibody incubation. Finally, cells were equilibrated in PBS, stained for DNA with DAPI (0.5 μg/ml), and mounted with Fluoromount G (Fisher Scientific, Newark, Del.).

To visualize lipid droplets, the cells were incubated with BODIPY 500/510 (Life Technology Corp. cat# B3824) at a final concentration of 1 μg/ml for 30 min at 37°C. the cells were then washed with PBS and applied for IFA as above.

### MTT [3-(4,5-Dimethylthiazol-2-yl)-2,5-Diphenyltetrazolium Bromide)] assay to evaluate cell viability

Cells were prepared in DMEM supplemented with 10% fetal bovine serum and seeded on 96-well plates. For the MTT assay, each drug was serially diluted by twofold dilutions. The highest concentration of 2-BP, 2-FPA, Cerulenin, or Palm B is 500 µM, 100 µM, 50 µM, or 50 µM respectively. The cell culture medium was replaced with the medium containing the diluted drugs. The 96-well plates were incubated at 37 °C for 24 or 48 hours. Cells were washed with phosphate-buffered saline (PBS) and incubated with complete MEM medium containing 0.5 mg/mL MTT reagent (sigma, lot#MKCD8033) for 2 hours. The medium was removed and 100 ul isopropanol was added into each well and incubated for 15 min. The absorbance of each sample was measured with a multi-plate reader spectrophotometer at 570 nm. Cell viability, proportional to the absorbance, was indicated by a relative reduction rate compared to the absorbance of no drug-treated cells (control). The viability of DMSO control cells was considered as 100%. Data are presented as mean ± standard error (SE) of at least three independent experiments.

### Immunoblot analysis

To determine the levels of cellular or viral proteins, the whole cell lysates were separated by SDS-PAGE in 7.5% polyacrylamide gels (10 to 20 µg loaded in each lane). The separated proteins in the gel were transferred to nitrocellulose membranes (Amersham Inc., Piscataway, NJ), and blocked with 5% nonfat milk for 60 min at room temperature. Membranes were incubated with a primary antibody, followed by an incubation with a horseradish peroxidase-coupled secondary antibody. For regular WB, we used secondary antibodies from Amersham. Detection was accomplished with enhanced chemiluminescence (Pierce, Rockford, IL), according to standard methods. Membranes were stripped with stripping buffer (100 mM β-mercaptoethanol, 2% SDS, 62.5 mM Tris-HCl, pH 6.8), washed with PBS-0.1% Tween 20, and re-probed with new primary antibodies to detect additional proteins. ImageJ was used for quantification of Western blot signals. All experiments were performed at least three times (n), and the values were presented as means ± SD; Statistical significance was analyzed by Student’s t test for pairwise comparisons and one way analysis of variance (ANOVA).

### Plaque formation unit (PFU) assay

The plaque formation unit (PFU) assay was used to determine viral titers, largely as described (24), but with a slight modification. Supernatants containing serially diluted virus particles were added to confluent 17CL-1 cell monolayers in 6-well plates. After adsorption for 2 h, the medium was removed, and the cells were washed twice with serum-free DMEM and overlaid with phenol-free DMEM containing 5% FCS, 0.5% low-melting point agarose (GIBCO), and 1% penicillin-streptomycin. Then, plaques were stained with Neutral red to enhance plaque visualization, and the plaque numbers were counted and calculated in 1 ml, which is pfu per ml. The mean pfu was determined after averaging from different dilutions.

### Immunofluorescent Microscopy

Cells were examined with an inverted Leica scanning system. Two or three channels were recorded simultaneously and/or sequentially and controlled for possible breakthrough between the fluorescein isothiocyanate and Texas Red signals and between the blue and red channels.

### APE (acyl-PEG exchange) assay

The cells were lysed with **4% (wt/vol) SDS** (Fischer) in **TEA buffer** [pH 7.3, 50 mM triethanolamine (TEA), 150 mM NaCl] containing 1x protease inhibitor mixture (Roche), 5 mM PMSF (Sigma), 5mM EDTA (Fischer), and 1,50 units/mL benzonase (EMD). The protein concentration of the cell lysate was then measured using a BCA assay (Thermo). Typically, 200 μg of total protein in 92.5 μL of lysis buffer was treated with 5 μL of 200 mM neutralized TCEP (Thermo) for final concentration of 10 mM TCEP for 30 min with nutation. NEM (Sigma), 2.5 μL from freshly made 1 M stock in ethanol, was added for a final concentration of 25 mM and incubated for 2 h at room temperature. Reductive alkylation of the proteins was then terminated by methanol-chloroform-H2O precipitation (4:1.5:3) with sequential addition of methanol (400 μL), chloroform (150 μL), and distilled H2O (300 μL) (all prechilled on ice). The reactions were then mixed by inversion and centrifuged (Centrifuge 5417R, Eppendorf) at 20,000 x g for 5 min at 4 °C. To pellet the precipitated proteins, the aqueous layer was removed, 1 mL of prechilled MeOH was added, the Eppendorf tube inverted several times and centrifuged at 20,000 x g for 3min at 4 °C. The supernatant was then decanted, and the protein pellet washed once more with 800 μL of prechilled MeOH, centrifuged again, and dried using a speed-vacuum (Centrivap Concentrator, Labconco). To ensure complete removal of NEM from the protein pellets, the samples were resuspended with 100 μL of TEA buffer containing 4% SDS, warmed to 37 °C for 10 min, briefly (∼5 s) sonicated (Ultrasonic Cleaner, VWR), and subjected to two additional rounds of methanol-chloroform-H2O precipitations, as described above. For hydroxylamine (NH2OH) cleavage and mPEG-maleimide alkylation, the protein pellet was resuspended in 30 μL TEA buffer containing 4% SDS, 4 mM EDTA and treated with 90 μL of 1 M neutralized NH2OH (J. T. Baker) dissolved in TEA buffer pH 7.3, containing 0.2% Triton X-100 (Fisher) to obtain a final concentration of 0.75 M NH2OH. Protease inhibitor mixture or PMSF should be omitted, as these reagents can interfere with the NH2OH reactivity. Control samples not treated with NH2OH were diluted in 90 μL TEA buffer with 0.2% Triton X-100. Samples were incubated at room temperature for 1 h with nutation. The samples were then subjected to methanol-chloroform-H2O precipitation, as described above, and resuspended in 30 μL TEA buffer containing 4% SDS, 4 mM EDTA, warmed to 37 °C for 10 min, and briefly (∼5 s) sonicated and treated with 90 μL TEA buffer with 0.2% Triton X-100 and 1.33 mM mPEG-Mal (5 or 10 kDa; Sigma) for a final concentration of 1 mM mPEG-Mal. Samples were incubated for 2 h at room temperature with nutation before a final methanol-chloroform-H2O precipitation. Dried protein pellets were resuspended in 50 μL 1Å∼ Laemmli buffer (Bio-Rad) and then heated for 5 min at 95 °C. Typically, 15 μL of the sample was loaded in 4–20% Criterion-TGX Stain Free polyacrylamide gels (Bio-Rad), separated by SDS/PAGE, and analyzed by Western blot.

### Statistical Analysis

All statistical significance was determined using either a paired or an un-paired student’s T-test depending on the experimental design. Statistical analysis of microscopy images was based on ImageJ quantification of randomly selected fields of view or cells (n>50) for each treatment or time point. Each study shown is representative of at least three independent experiments. The level of significance is denoted in each figure with P values.

## ACKNOWLEDGEMENT

The following reagent was obtained through BEI Resources, NIAID, and NIH: recombined Murine Coronavirus MHV-A59 with Enhanced Fluorescent Protein (eGFP), NR-53716; 17CL-1 (derived from NIH3T3), NR-53719; the monoclonal antibody against MHV-A59 nucleocapsid (N) protein, NR-45106. This study was supported by NIH/NIAID SC1AI112785 (Q.T.), and the National Institute on Minority Health and Health Disparities of the National Institutes of Health under Award Number G12MD007597 (Q.T.). The funders have no role in the study design, data collection, analysis, decision to publish, or preparation of the manuscript. The authors declare that they have no conflict of interest. This work was supported in part by Accelerating Excellence in Translational Science Pilot Grants G0812D05, NIH/NCI SC1CA200517 and 9 SC1 GM135050-05 to Y. Wu.

---

**Figure S1.**
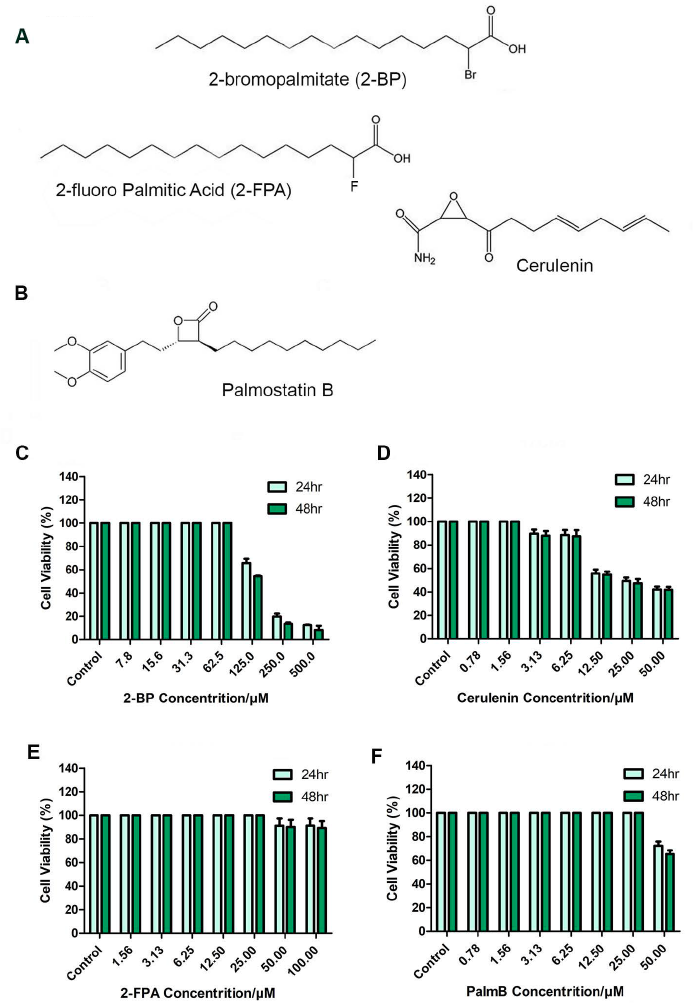
Palmitoylation inhibitors and enhancer and their toxicities to 17CL-1. **A.** Palmitoylation inhibitors: 2-BP, 2-FPA, and cerulenin. **B.** A palmitoylation enhancer: Palmostatin B. **C-F.** MTT [3-(4,5-Dimethylthiazol-2-yl)-2,5-Diphenyltetrazolium Bromide)] assay to evaluate cell viability after treatment of the palmitoylation inhibitors or enhancers. The cell viability (%) was compared to control (DMSO-treated) cells. Data were presented as mean ± standard error (SE) of at least three independent experiments.

**Figure S2.**
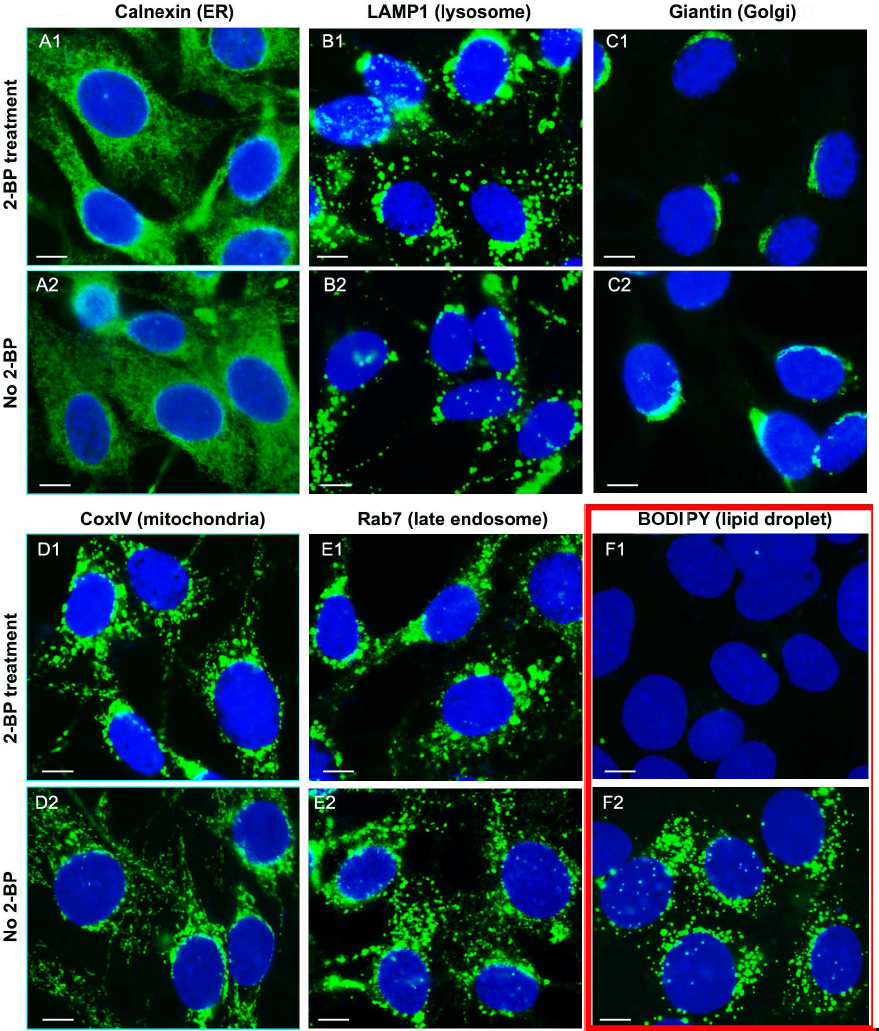
IFA to determine the effects of 2-BP on cellular organelles. After treatment of 25 uM 2-BP for 24 hours, the 17CL-1 cells were stained using the marker: calnexin for ER (A1-A2), LAMP1 for lysosome (B1-B2), Giantin for Golgi apparatus(C1-C2), CoxIV for mitochondria (D1-D2), Rab7 for late endosome (E1-E2), and BODIPY for lipid droplet (F1-F2). The picture was taken with a 100x lens. Scale bar: 10 μm.

**Figure S3.**
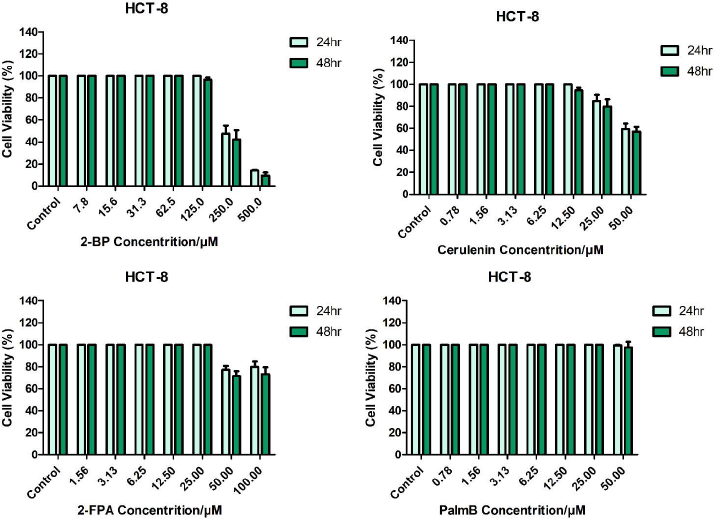
The cytotoxicities of 2-BP, 2-FPA, Cerulenin and palmostatin B to HCT-8 cells. Palmitoylation inhibitors: 2-BP, 2-FPA, and cerulenin and a palmitoylation enhancer: Palmostatin were added into HCT cells to to evaluate cell viability by MTT [3-(4,5-Dimethylthiazol-2-yl)-2,5-Diphenyltetrazolium Bromide)] assay. The cell viability (%) was compared to control (DMSO-treated) cells. Data were presented as mean ± standard error (SE) of at least three independent experiments.

**Figure S4.**
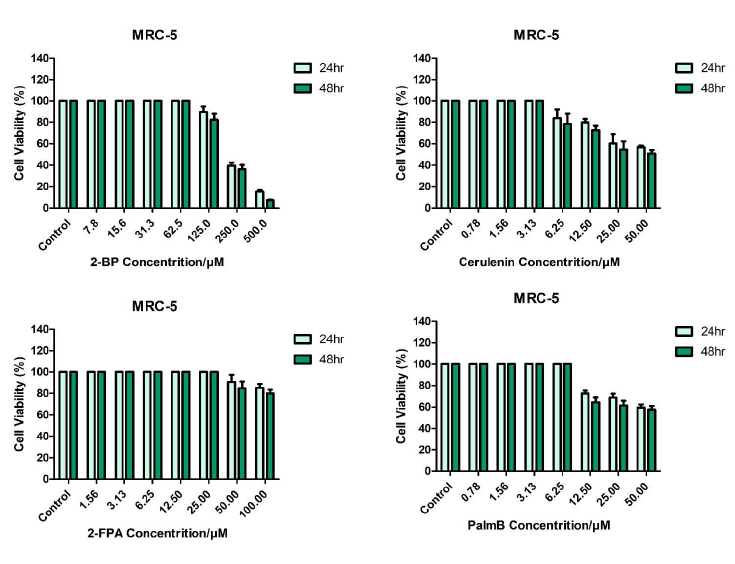
The cytotoxicities of 2-BP, 2-FPA, Cerulenin and palmostatin B to MRC-5 cells. The same as Figure S3 but in MRC-5 cells.

